# IL-33 promotes innate lymphoid cell-dependent IFN-γ production required for innate immunity to *Toxoplasma gondii*

**DOI:** 10.1101/2021.01.10.426122

**Authors:** Joseph T. Clark, David A. Christian, Jodi A. Gullicksrud, Joseph A. Perry, Jeongho Park, Maxime Jacquet, James C. Tarrant, Enrico Radaelli, Jonathan Silver, Christopher A. Hunter

## Abstract

IL-33 is an alarmin required for resistance to the parasite *Toxoplasma gondii*, but its role in innate resistance to this infection is unclear. *T. gondii* infection promotes increased stromal cell expression of IL-33 and levels of parasite replication correlate with IL-33 release. In response to infection, a subset of innate lymphoid cells (ILC) emerges composed of IL-33R^+^ NK cells and ILC1s. In Rag^-/-^ mice, where NK cells and ILC1 provide an innate mechanism of resistance to *T. gondii*, the loss of IL-33R reduced ILC responses and increased parasite replication. Furthermore, administration of IL-33 to Rag^-/-^ mice resulted in a marked decrease in parasite burden, increased production of IFN-γ and the recruitment and expansion of inflammatory monocytes associated with parasite control. These protective effects of exogenous IL-33 were dependent on endogenous IL-12p40 and the ability of IL-33 to enhance ILC production of IFN-γ. These results highlight that IL-33 synergizes with IL-12 to promote ILC-mediated resistance to *T. gondii*.

## Introduction

*Toxoplasma gondii* is an intracellular parasite of public health significance [1]–[3]. Resistance to this organism is initiated by dendritic cell production of IL-12, which promotes NK and T cell secretion of IFN-γ [4]–[6]. IFN-γ in turn induces multiple anti-microbial mechanisms, which include the activation of macrophages to express iNOS, which are required to limit parasite replication [7]–[12]. Previous studies have shown that mice deficient in the adapter molecule MyD88 have increased susceptibility to *T. gondii* associated with reduced production of IL-12 and IFN-γ [13],[14]. Since MyD88 is a major adapter required for Toll Like Receptor (TLR) signaling, this increased susceptibility is consistent with a role for TLR-mediated recognition of this pathogen [9],[10]. However, TLR1, TLR2, TLR4, TLR6, TLR9 and TLR11 are individually not required for early resistance to *T. gondii* [15]–[18], and there is a MyD88-independent mechanism of parasite recognition [19]–[21]. Moreover, administration of IL-12 to MyD88^-/-^ mice does not restore the ability to produce IFN-γ, and NK and T cell expression of MyD88 is required for optimal production of IFN-γ and resistance to *T. gondii* [22],[23]. Thus, MyD88 has a critical role in resistance to *T. gondii*, but the events that engage this adapter molecule are unclear.

Members of the IL-1 family of cytokines, including IL-1α/β, IL-18, and IL-33, utilize distinct receptor sub-units but share downstream signaling machinery that includes MyD88. These cytokines impact a wide range of immune cells, and influence many facets of the innate immune system [24]. Mice that lack T and B cells have helped define the impact of cytokines on innate mechanisms of immunity to a wide variety of pathogens [25]–[28]. For example, these models were important to identify the role of IL-12 in promoting NK cell production of IFN-γ required for resistance to *Listeria monocytogenes* and *T. gondii* [29]–[31]. It is now appreciated that NK cells and ILC1 populations are both relevant sources of IFN-γ that contribute to resistance to *T. gondii* [32],[33]. Although IL-1 and IL-18 synergize with IL-12 to promote NK cell production of IFN-γ [34]–[36], the role of endogenous IL-1 during toxoplasmosis is secondary to those of IL-12 [17],[34],[37] while endogenous IL-18 is not required for parasite control but rather contributes to the immune pathology that can accompany this infection [35],[38]–[41].

IL-33 is a cytokine that is constitutively expressed by endothelial and epithelial cells, and in current models the death of these cells leads to release of IL-33 that acts as a damage associated molecular pattern (DAMP) or alarmin to activate immune cell populations [42]–[45]. While there are open questions about whether this cytokine can also be secreted [46],[47], the rapid oxidation of IL-33 inactivates this cytokine and ensures that its activity is restricted to local sites of tissue damage [48]. IL-33 and the IL-33R ST2 are most prominently associated with amplification of TH2 CD4^+^ T cells, activation of ILC2 and resistance to helminths [42],[49]–[53], immune regulation by Treg cells [54],[55], and a number of metabolic and para-immune functions mediated by ILC2 and regulatory T cells [56],[57]. Consistent with its ability to promote TH2-type responses, IL-33 can antagonize inflammation mediated by TH1/TH17cells during experimental allergic encephalomyelitis (EAE) [58]–[60] and suppresses pathological TH1 responses during visceral Leishmaniasis [61]. However, during infection with LCMV or MCMV, IL-33 promotes the expansion of NK cells and T cells and their production of IFN-γ, and loss of the IL-33R results in a delay in viral clearance [62]–[64]. In contrast, perhaps one of the most strking phenotypes of mice that lack IL-33R is that they succumb to chronic toxoplasmosis, associated with reduced astrocyte responses required for protective T cell responses [65],[66]. Nevertheless, the acute stage of toxoplasmosis is associated with the ability of *T. gondii* to infect and lyse epithelial and endothelial cells [67]–[71], but whether these events lead to the release of IL-33 or if this affects the innate response to *Toxoplasma* is unknown. The present study reveals that parasite replication during acute toxoplasmosis is associated with release of IL-33, and Rag^-/-^ mice that lack the IL-33R have defects in ILC production of IFN-γ and impaired parasite control. Furthermore, administration of IL-33 to Rag^-/-^ mice enhanced ILC production of IFN-γ associated with the expansion of a population of Ly6c^hi^ CCR2^+^ inflammatory monocytes and a marked reduction in parasite burden.

Together, these results highlight that infection-induced release of IL-33 synergizes with IL-12 to promote ILC-mediated resistance to *T. gondii*.

## Materials and Methods

### Mice

C57BL/6NTac (Taconic #B6-F), Rag1^-/-^ (B6.129S7-Rag1^tm1Mom^/J) (Jackson #002216), and Rag2^-/-^Il2rg^-/-^ (Rag2^tm1Fwa^II2rg^tm1Wjl^) (Taconic #4111) mice were purchased from their respective vendors. IL-33^-/-^ (Il33^tm1.1Arte^)(Jackson #350163) mice were provided by MedImmune (now AstraZeneca). IL33^fl/fl^-eGFP (B6(129S4)-Il33^tm1.1Bryc^/J), originally generated by Paul Bryce were obtained locally from Dr. De’Broski Herbert, IL-33R^-/-^ *Il1rl1^-/-^* mice, originally derived by Andrew McKenzie (University of Cambridge) (16) and back-crossed to C57BL/6 by Peter Nigrovic (Harvard University), were provided by Edward Behrens at Children’s Hospital of Philadelphia. IL-33R^-/-^ Rag1^-/-^ mice were generated by crossing the knockouts described above. Mice were housed in a specific pathogen free environment at the University of Pennsylvania School of Veterinary Medicine and treated according to protocols approved by the Institutional Animal Care and Use Committee at the University. Male and Female (age 8–12 weeks at start of experiment) mice were used for all experiments.

### Parasites and Infection

The ME49 strain of *T. gondii* was maintained by serial passage in Swiss Webster mice and used to generate banks of chronically infected CBA/ca mice, which were a source of tissue cysts for these experiments. Pru-derived transgenic parasites and CPS parasites were maintained in cultured human fibroblasts in DMEM supplemented with 10% FBS. For CPS parasites, supplemental uracil was also added to media. Mice were infected intraperitoneally with 20 cysts (ME49), or 1×10^4^ tachyzoites (Pru), or 2×10^5^ tachyzoites (CPS). Soluble toxoplasma antigen was prepared from tachyzoites of the RH strain as described previously [72]. For quantitative PCR (qPCR), DNA was isolated from tissues using the DNEasy DNA isolation kit (Qiagen) followed by qPCR measuring the abundance of the *T. gondii* gene B1 using the primers 5’-TCTTTAAAGCGTTCGTGGTC-3’ (forward) and 5’-GGAACTGCATCCGTTCATGAG-3’ (reverse).

### Histology

For IHC detection of *T. gondii* and iNOS, tissues were fixed in 10% formalin solution and then paraffin embedded and sectioned. Sections were deparaffinized, rehydrated, Ag retrieved in 0.01 M sodium citrate buffer (pH 6.0), and endogenous peroxidase blocked by 0.3% H_2_O_2_ in PBS. After blocking with 2% normal goat serum, the sections were incubated either with *anti-Toxoplasma* Ab, anti-INOS Ab or isotype control. The sections were then incubated with biotinylated goat anti-rabbit IgG (Vector, Burlingame, CA), and ABC reagent was applied (Vectastain ABC Kit; Vector Labs). Then DAB substrate (Vector Labs) was used to visualize specific staining according to manufacturer’s instructions, and slides were counterstained with hematoxylin. To quantify parasite burden in the peritoneal exudate, 100,000 cells were used to prepare cytospins. Cells were methanol fixed and then stained with the Protocol Hema-3 Stain Set, and the ratio of infected cells to total cells in a field of view was calculated. For whole tissue mount immunofluorescence staining, omenta were harvested from mice and fixed in 1% PFA overnight at 4°C. After rinsing, tissue was blocked using 10% BSA, 0.5% normal rat serum (Invitrogen), and 1 μg/ml 2.4G2 (BD) in PBS for 1 hr at room temperature. Omenta were next incubated in PBS containing primary antibodies at 4°C for 3 days and subsequently rinsed with PBS overnight.

Immunofluorescence combining IL-33 (R&D AF3626) and CD45 (Biolegend 30-F11) antibodies was performed using the OPAL Automation Multiplex IHC Detection Kit (Akoya Biosciences, Catalog 160 #NEL830001KT) implemented onto a BOND Research Detection System (DS9455). All widefield images were obtained on a Leica DM6000 microscope using the Leica Imaging Suite software. Confocal images were acquired on a Leica STED 3X Super-resolution microscope. Image analysis was performed using FIJI and Imaris software packages.

### Generation of Lymphokine Activated Killer cells

Lymphokine Activated Killer cells (LAKs) were generated from Rag1^-/-^ bone marrow as described previously [73],[74]. Briefly, whole bone marrow was plated at 1M cells/mL in cRPMI + 400U/mL Proleukin human IL-2 (Peprotech). Fresh IL-2 was added every 3^rd^ day, and cells were used for experiments between days 7 to 10.

### Antibody and cytokine reagents

For in vitro assays, recombinant IL-33 was purchased from Peprotech (Cat #210-33 Rocky Hill, NJ). For in vivo treatment experiments, recombinant IL-33 (MedImmune), which was modified to be resistant to oxidation was used, as described previously[48]. IL-33 DuoSet ELISA was purchased from R&D Biosystems (Cat # DY3626, Minneapolis, MN). For flow cytometry the following combinations of antibodies were used: for analysis of NK cells: CD335 Nkp46 (29A1.4, eBioscience), NK-1.1 (PK136, Biolegend), IFN-γ (XMG1.2, eBioscience), CD200R1 (OX110, eBioscience), IL-33R (DJ8, MD Biosciences). For analysis of myeloid cells: CD11b (M1/70, eBioscience), CD11c (N418, Biolegend), Ly6c (HK1.4, Biolegend), Ly6g (1A8, Biolegend), CCR2 CD192 (SA203G11, Biolegend), CD64 FcgRI (X54-5/7.1, Biolegend), MHC II I-A/I-E (m5/114.15.2, eBioscience), iNOS (CXNFT, eBioscience), IL-33R (DJ8, MD Biosciences). Flow cytometry was performed on BD Fortessa and X-50 cytometers and data analysis was performed using Flowjo 9 and Flowjo 10 (Treestar), and Prism 7 and 8 (Graphpad). Uniform Manifold Approximation and Projection for Dimension Reduction (uMAP) analysis was performed using the uMAP plug-in (version: 1802.03426, 2018, ©2017, Leland McInness) for Flowjo (Version 10.53). The Euclidean distance function was utilized with a nearest neighbor score of 15, and a minimum distance rating of 0.5.

### Quantification and statistical analysis

All data are expressed as means ± SEM. For comparisons between two groups, the Student’s t-test was applied. For data with more than two data sets, one-way ANOVA coupled with Tukey’s multiple comparisons test was applied. Statistical details are indicated in figure legends.

## Results

### *Toxoplasma gondii* infection induces IL-33 upregulation and release

To determine the impact of *Toxoplasma* infection on IL-33 expression and secretion, C57BL/6 WT and Rag^-/-^ mice were infected intraperitoneally (i.p.) with the Me49 strain or the replication-deficient CPS strain of *T. gondii*, and the levels of IL-33 at local sites of infection and affected tissues assessed by ELISA. In the peritoneum of naïve WT and Rag^-/-^ mice the level of IL-33 was below the limit of detection (<10 pg/mL) (Fig 1A). Infection i.p. with 2×10^5^ tachyzoites of the non-replicating CPS strain did not cause parasite-induced host cell lysis and failed to elicit detectable IL-33 at 1 or 5 days post-infection (dpi) (data not shown). Infection of WT mice with 20 cysts of Me49 resulted in <1% infected cells in the peritoneum at 5 dpi, and IL-33 was not detected (Fig 1A). When Rag^-/-^ mice received the same challenge, there were 2-5% infected cells at 5 dpi, and low levels of IL-33 were detected (Fig 1A). To test if IL-33 levels were a function of parasite burden, WT and Rag^-/-^ mice were treated with anti-IFN-γ, which resulted in a 20-fold increase in parasite load (data not shown) and a 3-4-fold increase in the levels of IL-33 (Fig 1A). When these data sets were collated and quantity of parasite DNA plotted versus IL-33 concentration, there was a strong correlation between parasite burden and IL-33 levels (R = 0.7902) (Fig 1B). To determine if IL-33 was released in other tissues affected by *T. gondii*, tissue biopsies from the liver of WT mice at 10 dpi were prepared and placed in culture for 24 hours and IL-33 release measured. While basal levels of IL-33 were detected in tissues from naïve WT mice and mice injected with replication deficient CPS parasites, the biopsies from infected mice showed significantly elevated levels of IL-33 (Fig 1C). These results suggest that parasite replication and lysis of infected cells lead to IL-33 release.

**Figure 1:**
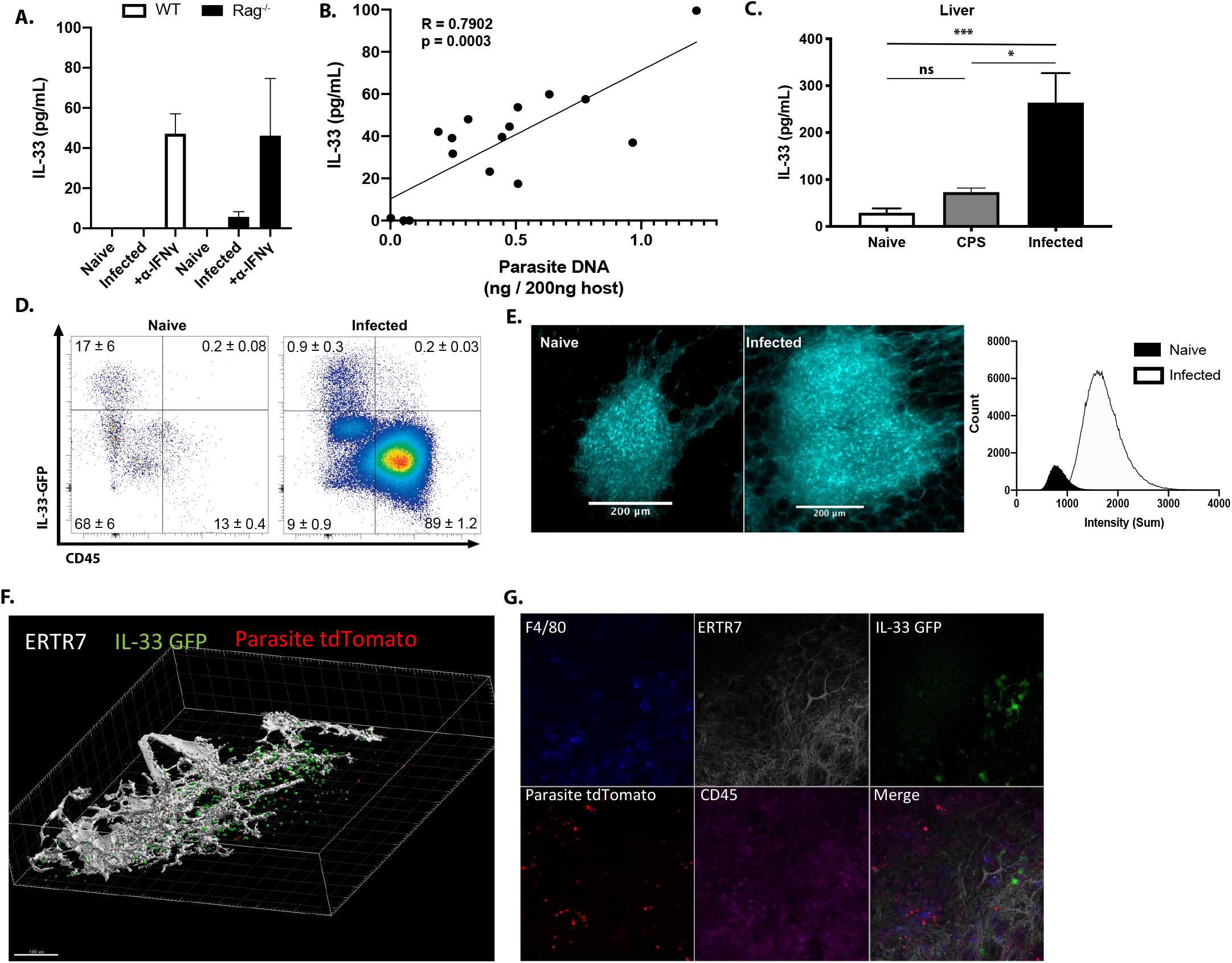
*Toxoplasma gondii* infection induces IL-33 expression and release. Mice were infected with *T. gondii* **(A)** after 7 days, free IL-33 in the peritoneal cavity was measured by ELISA. **(B)** Measurements of IL-33 from (A) were plotted against corresponding parasite burden and fit to a linear model. **(C)** 5mm punch biopsies of liver were placed in culture for 24 hours and IL-33 measured in supernatants by ELISA. 3 biopsies per mouse, 5 mice per group. **(D)** Cells from omenta of IL-33 GFP reporter mice were analyzed by flow cytometry at 3 days post infection. Cells shown are live singlets. Data are representative of 3 mice per group. **(E)** Whole mount omentum showing IL-33-GFP signal in milky spot. **(F)** 3D projection of milky spot showing stromal marker ERTR7 and IL-33 GFP signal. **(G)** Whole mount immunofluorescence of milky spot. NS, not significant (p>0.05); *p<0.05 and ***p<0.001 (one-way ANOVA with Tukey’s multiple comparisons test). Data are representative of or are pooled from three (A and B), or two (C, D, E, F and G) independent experiments (mean + s.e.m)

To identify the cellular source of IL-33 during infection, the IL-33-IRES-GFP mouse [75], a faithful reporter for IL-33 protein production (Supp Fig 1A), was utilized. The IL-33 reporter mice were infected i.p. with a fluorescent strain of *T. gondii* (Pru-tdTom), and the expression of IL-33-GFP in the omentum was examined by flow cytometry and IHC at 3 dpi. The omentum is an adipose tissue that contains Fat Associated Lymphoid Clusters (FALCs) which are one of the major sites for drainage from the peritoneum [76]–[78]. In naïve mice, the omentum contained a small population of CD45^+^ immune cells, most of which were IL-33-GFP^-^, whereas fibroblastic stromal cells (CD45^-^FSC^hi^ SSC^hi^ CD31^+/-^ PDPN^+/-^) were the main source of IL-33-GFP^+^ cells (Fig 1D and Supp Fig 1B). Upon infection, there was a marked expansion of the CD45^+^ population. A small population of CD45^+^ F4/80^+^ MHCII^+^ cells expressed IL-33 (data not shown) but the majority of IL-33-GFP^+^ cells remained fibroblastic stromal cells. The ability to detect infected cells based on parasite expression of tdTomato revealed that infected cells were not associated with IL-33 expression (Supp. Fig 1C). Similarly, at 7 days post infection the use of flow cytometry and immunofluorescence revealed that CD45^-^ cells were the dominant source of IL-33 in the spleen, lung, and liver (Supp Fig 1D).

To understand the spatial organization of the IL-33-GFP^+^ cells, the omentum was used for whole tissue mount immunofluorescence. In uninfected mice, consistent with the analysis above, IL-33 was constitutively expressed by non-hematopoietic CD45^-^cells with fibroblastic morphology distributed throughout the FALCS. At 3 dpi there was a marked increase in the size of the FALC, and an approximate 3-fold increase in number of IL-33^+^ cells (Fig 1E). These images are max projection views that illustrate the size of FALCs, but quantification of the intensity of fluorescence highlighted the 10-fold increase in the expression of IL-33-GFP (Fig 1E) associated with ERTR7^+^ fibroblastic reticular cells (Fig 1F). Imaging revealed that areas of parasite replication were inversely correlated with the presence of IL-33-GFP expression (Fig 1G). Together, these data establish that in vivo infection with *T. gondii* leads to the release of IL-33 by stromal cells that correlates with levels of parasite replication.

### ILC responses to IL-33

To identify the cell populations that could respond to the local release of IL-33 during this infection, a UMAP analysis was used to provide an unbiased comparison of the changes in IL-33R expression in the peritoneum of naïve and infected Rag^-/-^ mice (Fig 2A). In naïve mice, IL-33R was expressed by peritoneal macrophages (CD64^+^CD11b^+^MHCII^+/-^) and a small population of ILC2 (Lin^-^ Nkp46^-^) when compared with IL-33R^-/-^ controls (Fig 2A). By 5 dpi there was a marked change in the cellular composition of the peritoneum with a loss of the MHCII^-^ macrophage and ILC2 populations but a prominent monocyte and neutrophil infiltration and the expansion of NKp46^+^ NK cells and ILC1s. While there were low levels of IL-33R expressed by MHCII^hi^ CD64^+^ cells, the highest levels of IL-33R were observed on NKp46^+^ cells. Further validation revealed that in naïve WT, IL-33R^-/-^, and IL-33^-/-^ mice, IL-33R expression was not detected on peritoneal or splenic NK cells, but IL-33R was observed on a subset (~20%) of NK cells by 5 dpi (Fig 2B). Furthermore, NK cells from infected IL-33^-/-^ mice still upregulated IL-33R, indicating that IL-33 signaling is not required for this process. Thus, infection with *T. gondii* leads to the emergence of populations of NK cells and ILC1s that express the IL-33R.

**Figure 2:**
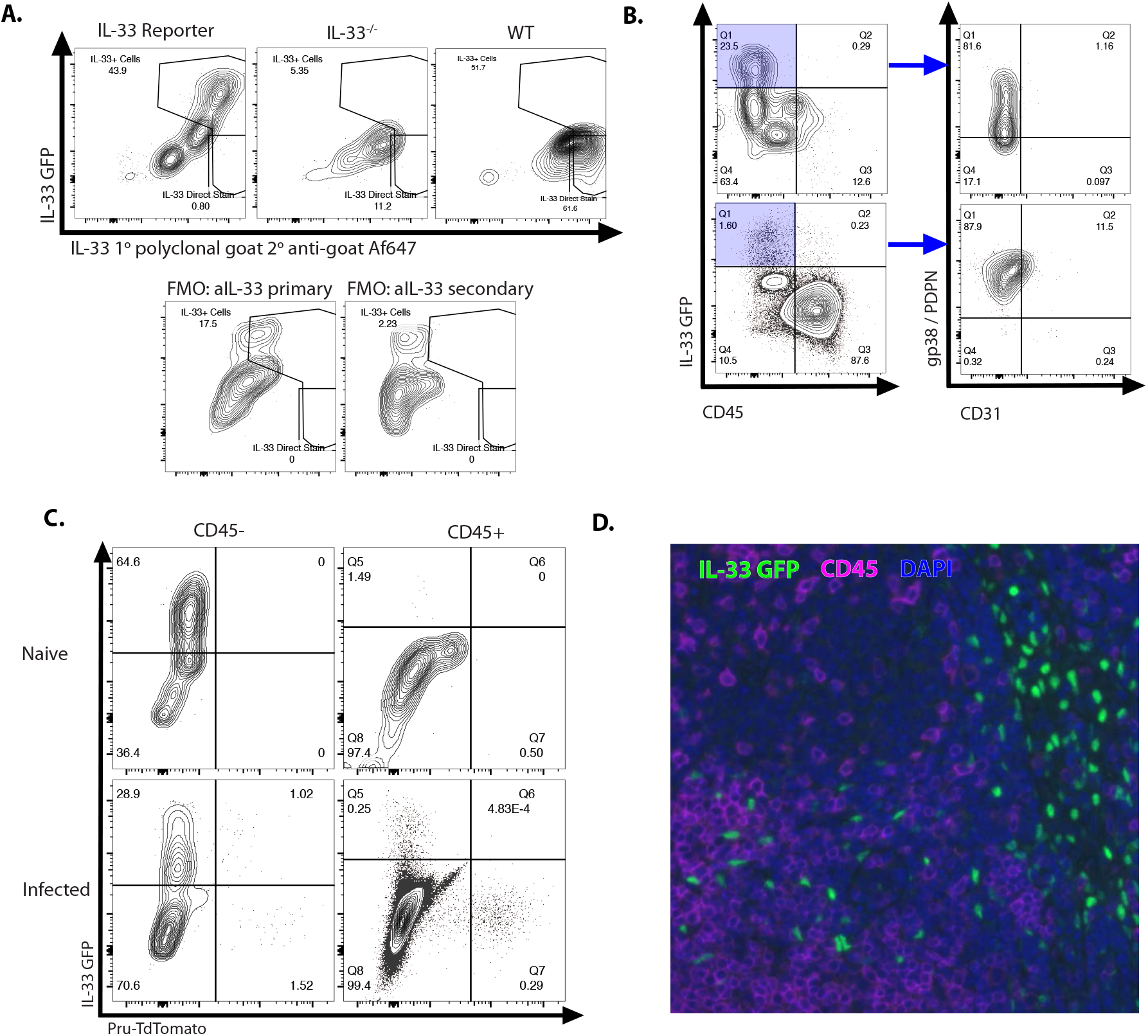
Infection sensitizes NK and ILC to IL-33. **(A)** UMAP analysis of peritoneal exudate cells from naïve or 7 dpi mice, with heatmap for IL-33R expression. Data compiled from 4 mice per group. **(B)** Flow cytometry from peritoneal cells showing IL-33R staining on NKp46+ cells. Data are representative of 3-4 mice per group. **(C)** Flow cytometry of LAKs showing composition of population based on cytokine stimulation condition. Population shown is pre-gated on live singlets. **(D)** Intracellular cytokine staining of LAKs after 24-hour cytokine stimulation and 4 hour incubation with Brefeldin A. Data are representative of 3 independent experiments. NS, not significant (p>0.05) (student’s t-test); Data are representative of or are pooled from three independent experiments (A-D).

Next, a population of IL-2 induced LAKs (generated from the bone marrow of Rag^-/-^ mice) were utilized to compare the impact of IL-33 (and its relative IL-18) alone or in combination with IL-12 on ILCs. Phenotyping of these LAK cultures revealed that they contained ILC1s (NKp46^+^ CD200R1^+^), ILC2s (NKp46^-^ CD200R1^+^) and NK cells (NKp46^+^ CD200R1^-^) (Fig 2C). Upon withdrawal of IL-2, the addition of IL-33 preferentially stimulated the proliferation of CD200R1^+^ ILC2s, while IL-18 stimulated NK cell proliferation (Fig 2C). IL-12 alone did not induce the expansion of a specific cell type, but when combined with IL-33 maintained the heterogeneity of the LAK population and IL-12 plus IL-18 resulted in a modest increase in the proportion of NK cells compared to IL-18 alone. Testing of the ability of IL-33 to stimulate LAKs to produce IFN-γ showed that IL-33 alone did not stimulate LAKs to produce IFN-γ but did act in synergy with IL-12 to enhance the production of IFN-γ in the NKp46^+^ populations (Fig 2D). Similar results were observed when splenocytes from Rag^-/-^ mice were used (Fig 2D), indicating that these effects of IL-33 on NK cells were not dependent on pre-activation with IL-2.

These observations are consistent with previous reports on the ability of IL-33 to promote ILC2 activity [26],[79], but demonstrate that in the presence of IL-12, IL-33 is a potent inducer of IFN-γ.

### Endogenous IL-33 is required for innate resistance to *T. gondii*

To directly test the role of endogenous IL-33 in innate resistance to *T. gondii*, Rag^-/-^ mice that lacked the IL-33R (Rag^-/-^IL-33R^-/-^ mice) were generated and infected i.p. with *T. gondii*. Compared to Rag^-/-^ mice, at 7 dpi the Rag^-/-^IL-33R^-/-^ mice showed an increased parasite burden based on the frequency of infected cells in the peritoneum (Fig 3A) and quantitation of parasite DNA in the peritoneum and liver (Fig 3B). Serum analysis of infected mice revealed comparable levels of IL-12p40 in Rag^-/-^ and Rag^-/-^IL-33R^-/-^ mice, but IFN-γ was severely compromised in the absence of IL-33R (Fig 3C). At this time point, Rag^-/-^ mice had a marked expansion in ILC1s and NK cells in the liver that was reduced in the absence of the IL-33R (Fig 3D). Consistent with decreased production of IFN-γ, fewer Ly6c^hi^ monocytes were recruited to the liver in the Rag^-/-^IL-33R^-/-^ mice, and these Ly6C^hi^ monocytes expressed lower levels of iNOS (Fig 3E). Analysis of the Ly6c^hi^ population in the peritoneum at 7 dpi after infection with Pru-tdTom showed that a proportion of infected and uninfected cells express iNOS in the Rag^-/-^ mice, but iNOS levels were markedly reduced in the Rag^-/-^IL-33R^-/-^ mice (Fig 3F). These data sets establish that endogenous IL-33 is required for optimal production of innate IFN-γ and the recruitment of monocyte populations that express anti-microbial effector mechanisms required for resistance to *T. gondii*.

**Figure 3:**
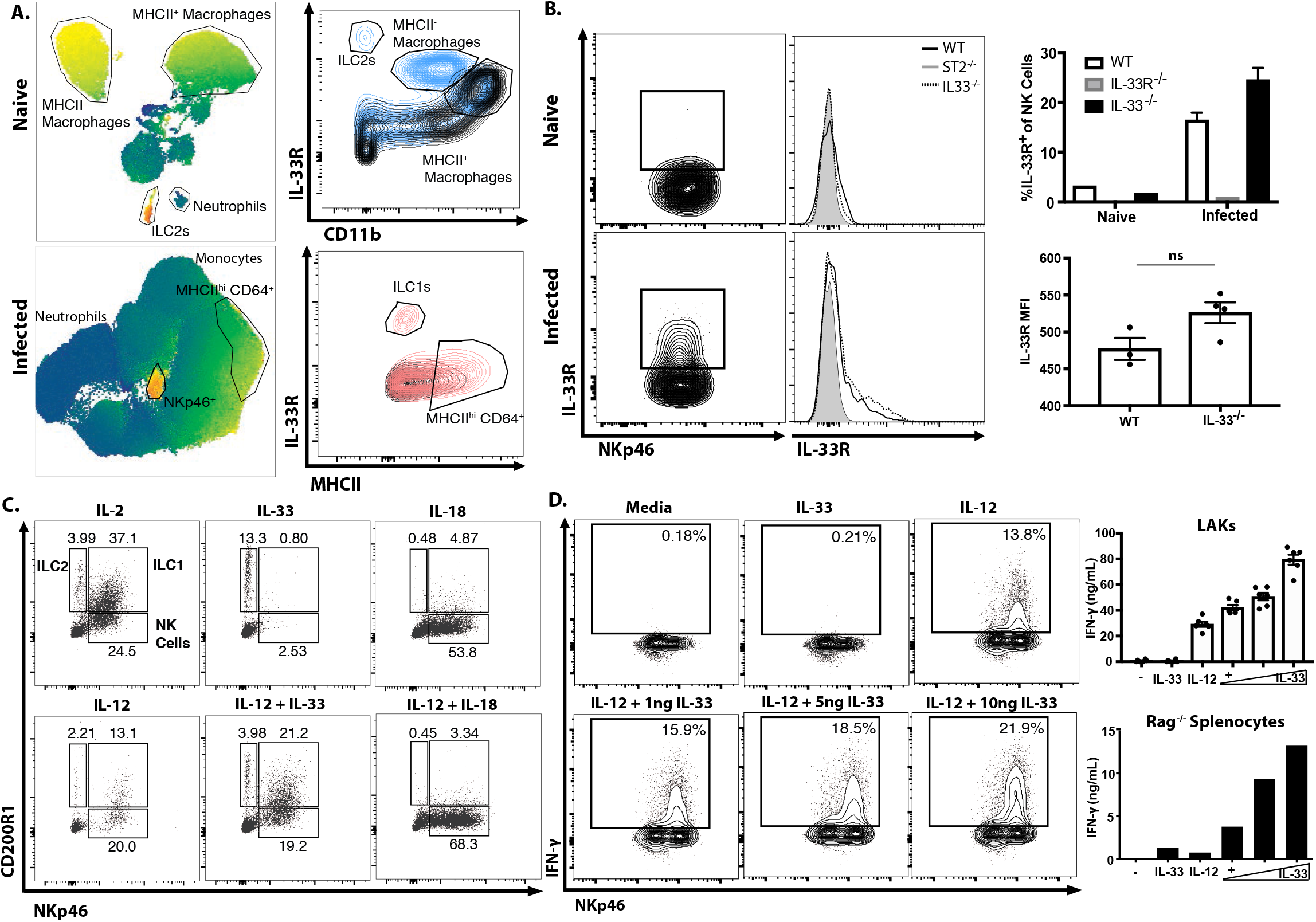
Endogenous IL-33 promotes the anti-parasitic immune response. **(A)** Cytospins of peritoneal exudate cells at 7 dpi. **(B)** qPCR for parasite DNA from indicated tissues. **(C)** Serum cytokines measured by ELISA at 7 dpi. Representative of 4-5 mice per group **(D)** Flow cytometric analysis and quantification of liver innate lymphoid cells. Populations shown are pre-gated on live singlets that are MHCII^-^. **(E)** Quantification of inflammatory monocytes (CD11b+ CD64+ Ly6g-) in livers of infected mice at 7 dpi. **(F)** Intracellular iNOS staining from monocytes in (E). NS, not significant (p>0.05); *p<0.05, **p<0.01, ***p<0.001, and ****p<0.0001 (student’s t-test). Data are representative of 3 independent experiments.

### IL-33 treatment boosts IL-12 and IFN-γ dependent immunity

Based on the ability of IL-33 to stimulate IL-12-dependent IFN-γ production in ILC1s and NK cells, a recombinant version of IL-33, resistant to oxidation which has a 30 fold increase in efficacy [36],[48], was utilized to determine if exogenous IL-33 could be used to enhance innate resistance to *T. gondii*.

Beginning at 1 dpi, IL-33 was administered i.p. every two days until 7 dpi, which resulted in a dose-dependent reduction in the frequency of infected cells at the site of infection and a decrease in parasite DNA in multiple tissues (Fig 4A). Analysis of cytospins of PECs revealed that treatment of infected Rag^-/-^ mice with IL-33 resulted in the emergence of a highly activated monocyte population (Fig 4B). These inflammatory monocytes were larger (higher FSC) and more granular (higher SSC) (Fig 4C). Furthermore, these cells were characterized by their expression of CD11b, CD11c, Ly6c, CCR2, and MHCII (Fig 4C). IL-33 treatment also resulted in increased recruitment of Ly6c^hi^ CCR2^+^ inflammatory monocytes to the liver and lungs by 7 dpi, and these monocytes had enhanced iNOS and IL-33R expression (Fig 4D). Histological analysis of the liver confirmed that IL-33 treatment resulted in increased cellular infiltration and expression of iNOS (Fig 4E, black arrows). Importantly, these changes induced by IL-33 treatment were associated with decreased necrotic foci that were frequent in infected Rag^-/-^ mice (blue arrow). These results correlate the protective effects of IL-33 treatment with an increase in macrophage and monocytes responses required for the control of *T. gondii*.

**Figure 4:**
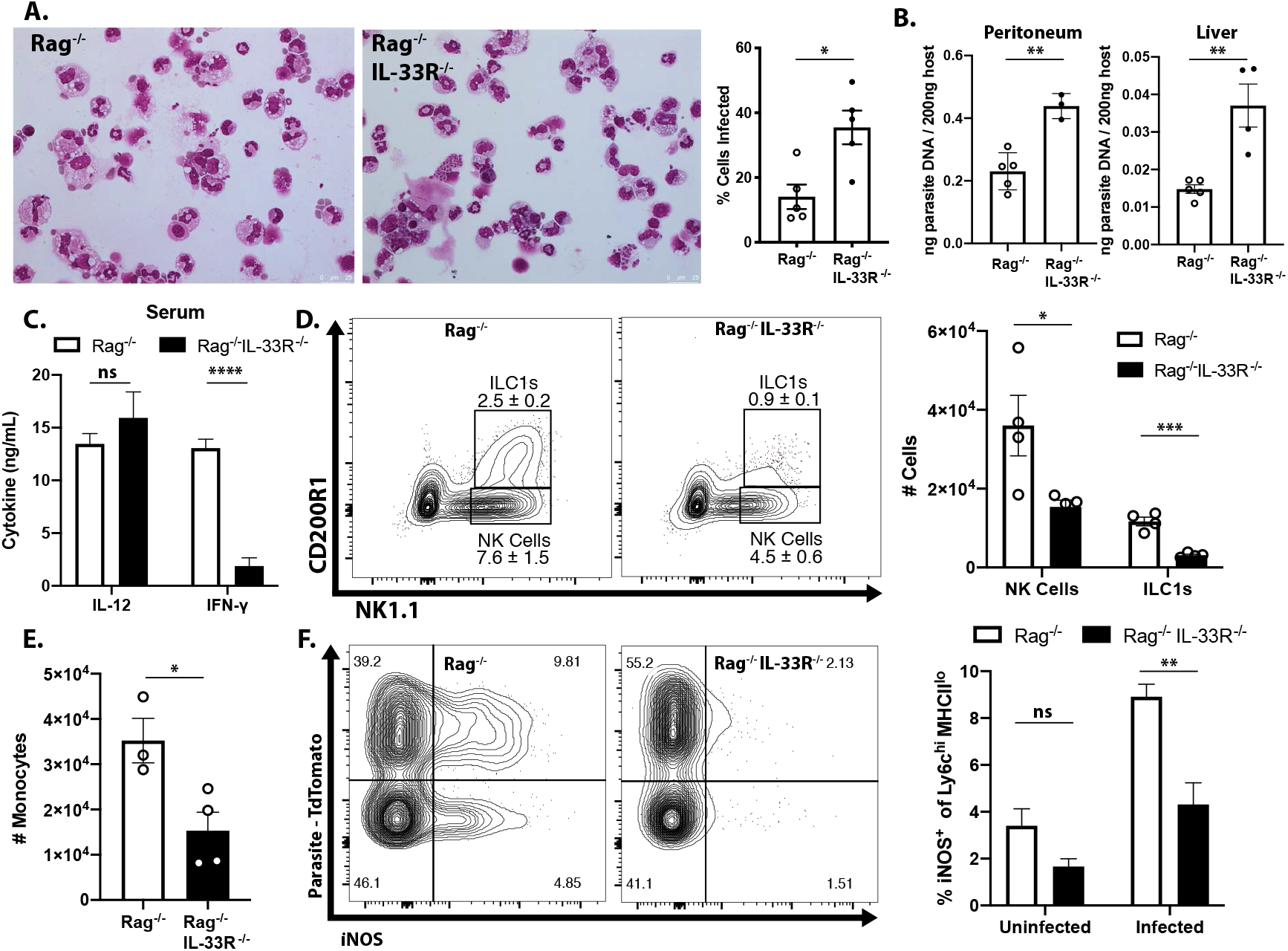
IL-33 treatment boosts IL-12 and IFN-γ dependent immunity. **(A)** Quantification of infected cell frequencies in cytospins at 7 dpi and qPCR for parasite DNA in indicated tissues. **(B)** Representative cytospins from peritoneal lavage at 7 dpi. Data are representative of 4-6 mice per group and 5 independent experiments. **(C)** Flow cytometric analysis of inflammatory monocytes in the peritoneal exudate at 7 dpi. Populations shown are pre-gated on live Ly6g^-^ singlets. Ly6c^hi^ CCR2^+^ cells are highlighted in black. **(D)** Representative analysis of Ly6c^+^ CCR2^+^ cells at 7 dpi in the liver and quantification of monocyte numbers and iNOS staining. **(E)** Histology of liver at 7 dpi, H&E showing infiltration of immune cells (left) and DAB iNOS staining (right). *p<0.05, **p<0.01, and ***p<0.001 (student’s t-test). Data are representative of or are pooled from three independent experiments.

To determine whether the protective effects of exogenous IL-33 depended on the ability of IL-12 to promote ILC production of IFN-γ, infected Rag^-/-^ mice were treated with IL-33 in combination with either anti-IL-12p40 or anti-IFN-γ neutralizing antibodies. Additionally, Rag^-/-^ γc^-/-^ mice, which lack ILC, were treated with PBS or IL-33. Blockade of either IL-12 or IFN-γ entirely abrogated the protective effects of IL-33 treatment as measured by the frequency of infected cells in the peritoneum at 7 dpi (Fig 5A). Rag^-/-^ γc^-/-^ mice were more susceptible than Rag^-/-^ mice, as expected, and IL-33 treatment did not affect parasite burden in the peritoneum. IFN-γ levels at the site of infection were increased by IL-33 treatment in ILC-sufficient Rag^-/-^ animals, but were unaffected in Rag^-/-^γc^-/-^ animals (Fig 5B). The expansion of Ly6c^hi^ CCR2^+^ monocytes associated with protection was also dependent on these factors, as cytokine blockade or absence of innate lymphoid cells effectively eliminated these cells (Fig 5C and Supp Fig 2). These results emphasize that the protective effects of IL-33 are dependent on IL-12 and ILC cytokine production and consequent recruitment of inflammatory monocytes.

**Figure 5:**
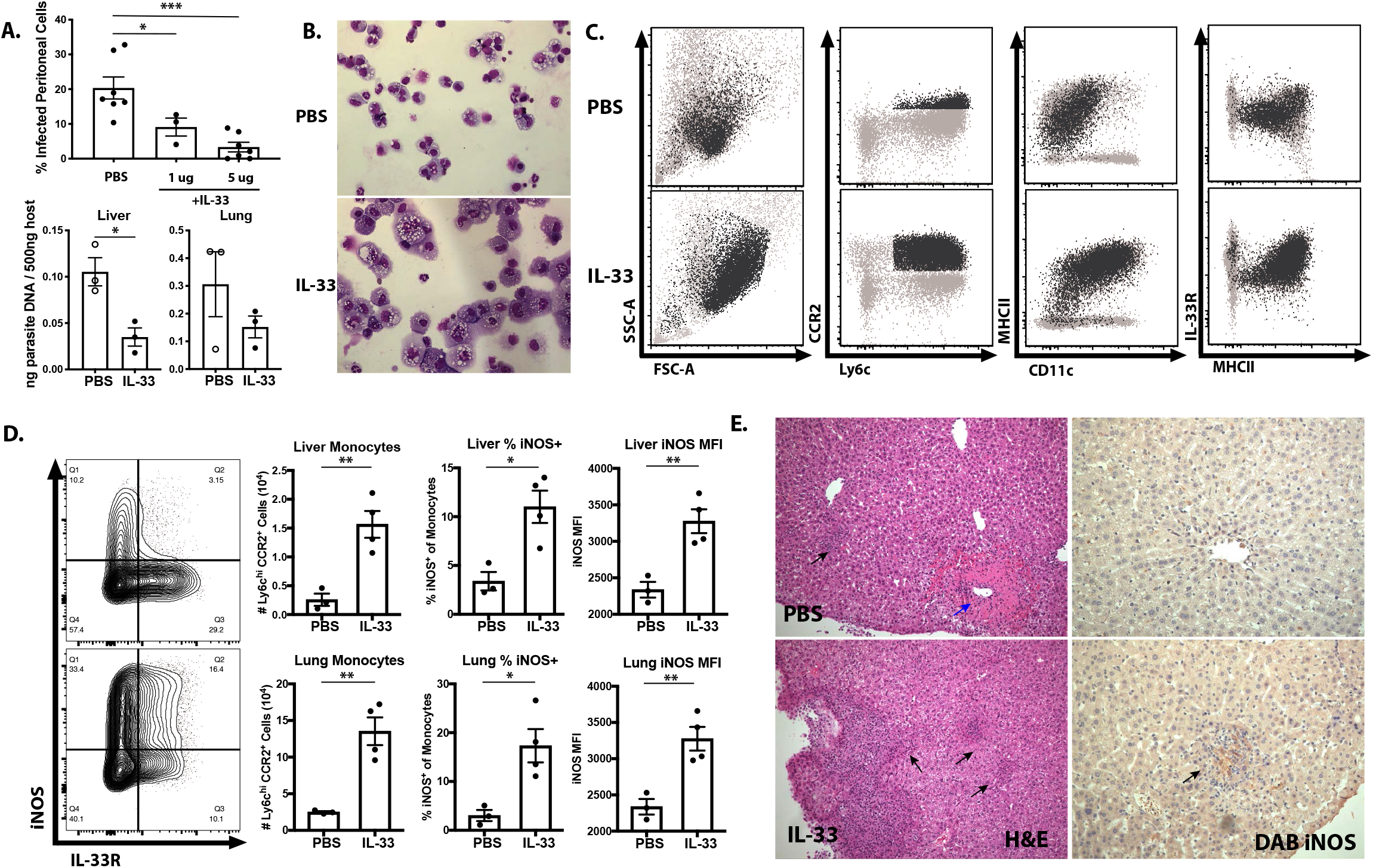
Protective effect of IL-33 is dependent on IL-12, IFN-γ, and ILC. **(A)** Quantification of cytospins from peritoneal exudate cells at 7 dpi. **(B)** Quantification of IFN-γ in peritoneal lavage at 7 dpi. **(C)** Flow cytometric analysis of inflammatory monocytes in peritoneum at 7 dpi. Data are representative of 5 mice per group and 2 independent experiments.

## Discussion

Previous studies have identified a central role for IL-12 in innate and adaptive production of IFN-γ required for control of *T. gondii* [4],[5],[80]–[83] but other cytokines and costimulatory pathways potentiate the effects of IL-12 on NK cells. In particular, IL-1 and IL-18 can amplify NK cell production of IFN-γ but evidence that endogenous IL-1 and IL-18 are critical for control of *T. gondii* in this model is limited. Thus, while IL-1 or IL-18 contribute to the development of infection-induced, microbiome-dependent immune mediated pathology in the gut, there is limited evidence that loss of IL-1 or IL-18 leads to increased parasite replication [17],[22],[37],[39],[84]–[87]. Indeed, early studies showed that neutralization of endogenous IL-18 did not affect levels of parasite replication, and that in SCID mice treated with IL-12 the effects of IL-1R blockade were modest and these treated mice were still more resistant than untreated SCID mice [34]. It is relevant to note that recent work has highlighted a cell intrinsic role for MyD88 in NK cells to help control *T. gondii* [23], but neither IL-1 blockade or the use of IL-1R^-/-^ and IL-18^-/-^ mice, replicates the susceptibility of MyD88^-/-^ mice [22],[17]. In the studies presented here, the reduced NK and ILC responses observed in Rag^-/-^IL-33R^-/-^ mice suggest that the ability of IL-33 (rather than IL-1 or IL-18) to amplify the IL-12-mediated innate response to acute toxoplasmosis helps explain the role for MyD88 in innate resistance to *T. gondii*. It is increasingly appreciated that in addition to NK cells, tissue resident ILC1s are an early source of IFN-γ in the innate response to *T. gondii* [32],[33], but the differential programming of these cells, including their responsiveness to the IL-1 family member cytokines, is still being described. While IL-12 is central to NK and ILC1 production of IFN-γ, there are other stimuli that can potentiate this pathway [88]. Certainly in vitro, in the presence of IL-12, IL-33 can promote NK cells and ILC1 IFN-γ production. While IL-18 is a more potent stimulator of IFN-γ in LAK cultures, the ability of IL-33 to promote NK and ILC1 responses, combined with the defects seen in these populations in IL-33R^-/-^ mice, suggest that of all the IL-1 family members IL-33 is uniquely important for innate lymphoid cell responses to *Toxoplasma*.

Because the signaling pathways for the IL-1 family members converge on MyD88-dependent activation of NF-kB, differences in expression patterns and tissue localization are likely to dictate the relative importance of each cytokine. The release of IL-1 and IL-18 is typically considered to be downstream of inflammasome mediated caspase activation and processing of pro-forms of these cytokines [89]. While there is evidence that pro-IL-1 and pro-IL-18 are produced during toxoplasmosis [31],[90]) several studies have concluded that *T. gondii* does not readily activate inflammasomes and there is evidence that *Toxoplasma* suppresses inflammasome activity [40], [91],[92]. However, there is a report that the inflammasome sensors NLRP1 and NLRP3 are required for protective immunity to *T. gondii* [41]. A possible explanation for this discrepancy is that some inflammasome components are not just microbial sensors but have additional functions that include a role for Caspase 8 in the activation of the c-Rel transcription factor required for expression of IL-12 and resistance to *T. gondii* [93]. To date, no murine sensor of *Toxoplasma* or parasite ligand has been identified that directly activates inflammasomes, although there is evidence for sensor-independent routes for inflammasome activation [94]. In contrast to the complex events that lead to the production and processing of IL-1 and IL-18, IL-33 is expressed constitutively by epithelial and endothelial cells at barrier sites and stored in the nucleus and therefore may be resistant to parasite mechanisms of immune evasion and suppression that target host cell transcription. The release of IL-33 can occur as a consequence of tissue damage associated with allergic inflammation or viral infection [63],[50] and during toxoplasmosis IL-33 levels correlated with levels of parasite replication. Thus, even though there may be non-canonical pathways for IL-33 release [46],[47],[56],[95],[96] it seems likely that these levels are a consequence of parasite-mediated lysis of infected cells.

Treatment of infected mice with exogenous IL-33 confirmed the protective effects of IL-33 and highlighted the impact on the recruitment of inflammatory monocytes to sites of infection and the subsequent upregulation of iNOS, a process required for control of *T. gondii* [7]–[10]. IL-33 drives ILC-cell dependent recruitment of CCR2^+^ inflammatory monocytes which resemble the TipDCs (TNF and iNOS-producing dendritic cells) previously recognized to be important for control of infection [57],[58]. This agrees with findings in an allergy model that described a role for IL-33 in the CCR2-dependent recruitment of inflammatory monocytes [97]. IL-33 may also act directly on these monocytes, as expression of the IL-33R was observed on monocytes in these studies, and it has been reported that IL-33 can directly enhance monocyte production of iNOS [98]. It is important to note that endogenous IL-33 is susceptible to rapid inactivation via oxidation in the extracellular space, which restricts its effects spatially and temporally. However, the recombinant IL-33 used in these studies was engineered to resist oxidation and it is possible that this treatment approach may have wider activities on hematopoiesis than IL-33 produced at sites of inflammation.

While IL-33 is most prominently linked to the regulation of Th2 type responses, there are reports that highlight the context dependent role that IL-33 plays in TH1 responses. In models of Leishmaniasis and cerebral malaria, IL-33 contributes to T-cell dependent immune pathology in the skin and brain, respectively [61],[99]. However, with the viral pathogens MCMV and LCMV, IL-33 contributes to NK and T cell expansion, and in its absence there is a delay in viral clearance, but in neither case is IL-33 essential for protective immunity [37],[62]. Indeed, during intracerebral LCMV infection IL-33 contributes to the development of lethal immune pathology [63], whereas for mice chronically infected with *T. gondii* the loss of IL-33 results in increased parasite burden [66]. More recent studies have highlighted that IL-33 promotes astrocyte responses that promote T cell responses required for control of *T. gondii* in the CNS [65]. Nevertheless, the data presented here establish that the ability of IL-33 to amplify ILC responses and their production of IFN-γ plays a protective role in the acute innate response to *Toxoplasma*. These results are consistent with a model in which IL-33 has a protective rather than pathological role in the immune response to *T. gondii*.

---

**Supplemental Figure 1:**
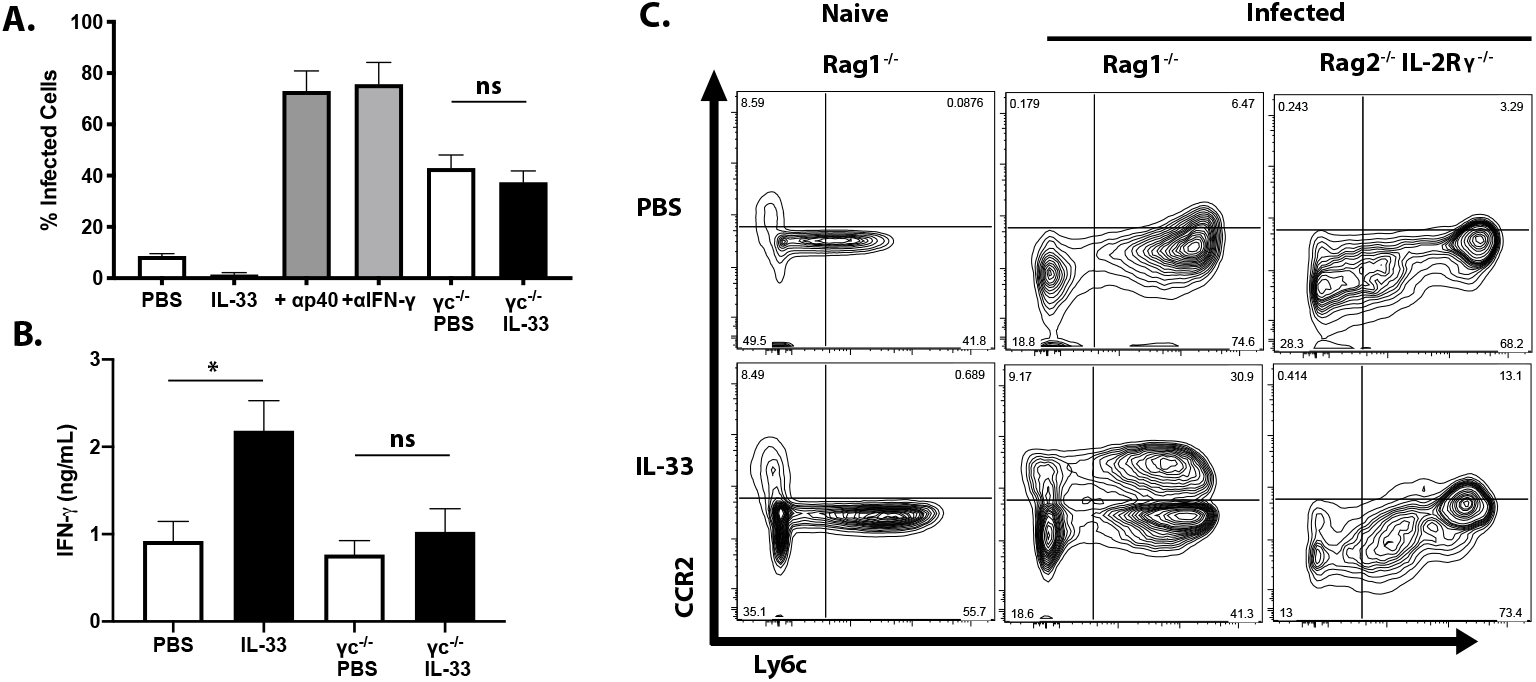
(A-C) Intracellular staining of omenta from infected mice. Data are representative of 3 mice per group. (D) Immunofluorescence of spleen sections from naïve mouse.

**Supplemental Figure 2:**
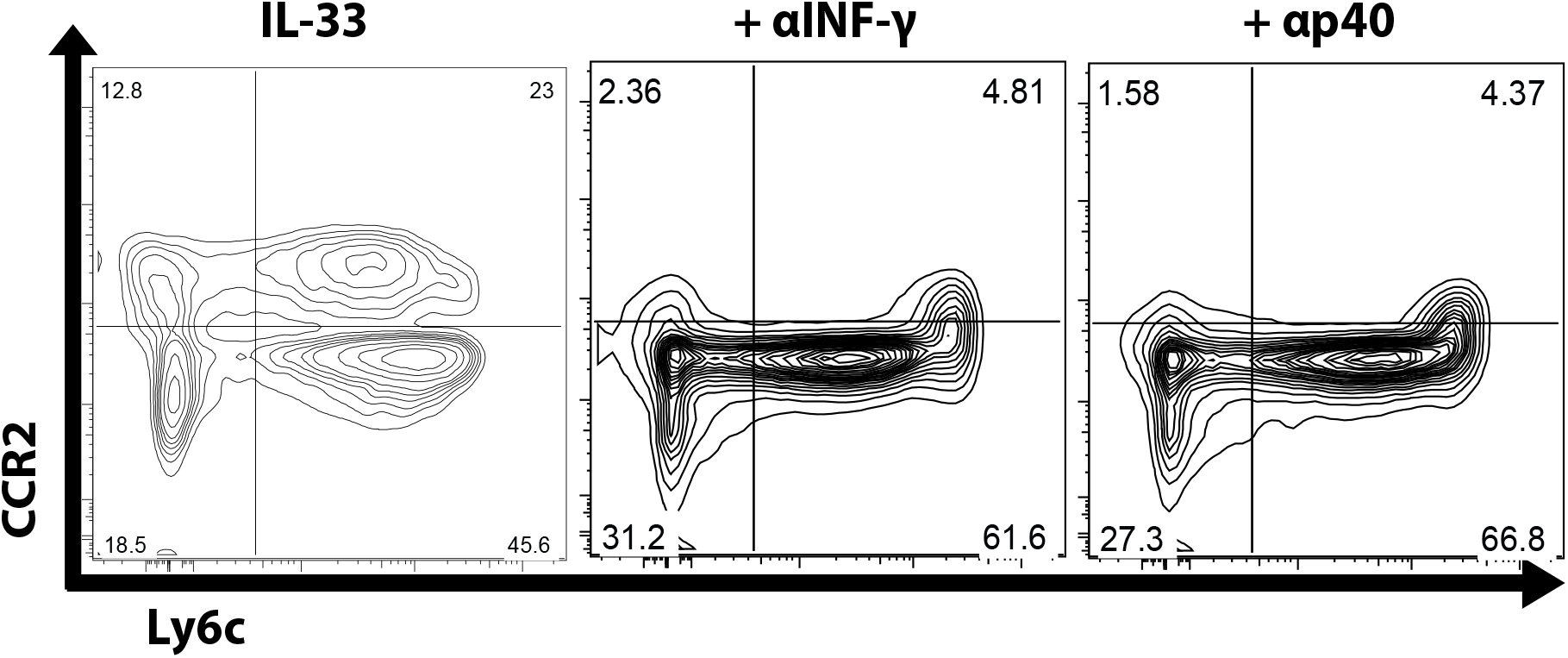
(A) Flow analysis of inflammatory monocyte recruitment in IFN-γ and IL-12p40 blockaded mice

## References

1. Montoya, J. G. & Liesenfeld, O. Toxoplasmosis. in Lancet 363, 1965–1976 (Elsevier, 2004).

2. Weiss, L. M. & Dubey, J. P. Toxoplasmosis: A history of clinical observations. Int. J. Parasitol. 39, 895–901 (2009).

3. Israelski, D. M. & Remington, J. S. Toxoplasmic encephalitis in patients with AIDS. Infect. Dis. Clin. North Am. 2, 429–45 (1988).

4. Khan, I. A., Matsuura, T. & Kasper, L. H. Interleukin-12 enhances murine survival against acute toxoplasmosis. Infect. Immun. 62, 1639–42 (1994).

5. Hunter, C. A., Subauste, C. S., Van Cleave, V. H. & Remington, J. S. Production of gamma interferon by natural killer cells from Toxoplasma gondii-infected SCID mice: regulation by interleukin-10, interleukin-12, and tumor necrosis factor alpha. Infect. Immun. 62, 2818–24. (1994).

6. Yap, G. S. & Sher, A. Cell-mediated immunity to Toxoplasma gondii: Initiation, regulation and effector function. in Immunobiology 201, 240–247 (Elsevier GmbH, 1999).

7. Yap, G. S. & Sher, A. Effector cells of both nonhemopoietic and hemopoietic origin are required for interferon (IFN)-γ- and tumor necrosis factor (TNF)-α-dependent host resistance to the intracellular pathogen, Toxoplasma gondii. J. Exp. Med. 189, 1083–1091 (1999).

8. Dunay, I. R., Fuchs, A. & Sibley, L. D. Inflammatory monocytes but not neutrophils are necessary to control infection with Toxoplasma gondii in mice. Infect. Immun. 78, 1564–70 (2010).

9. Serbina, N. V, Salazar-Mather, T. P., Biron, C. A., Kuziel, W. A. & Pamer, E. G. TNF/iNOS-roducing dendritic cells mediate innate immune defense against bacterial infection. Immunity (2003).

10. Scharton-Kersten, T. M., Yap, G., Magram, J. & Sher, A. Inducible nitric oxide is essential for host control of persistent but not acute infection with the intracellular pathogen Toxoplasma gondii. J. Exp. Med. 185, 1261–1273 (1997).

11. Wang, S. et al. Infection-induced intestinal dysbiosis is mediated by macrophage activation and nitrate production. MBio 10, (2019).

12. Chen, L. et al. The Toxoplasma gondii virulence factor ROP16 acts in cis and trans, and suppresses T cell responses. J. Exp. Med. 217, (2020).

13. Scanga, C. A. et al. Cutting edge: MyD88 is required for resistance to Toxoplasma gondii infection and regulates parasite-induced IL-12 production by dendritic cells. J. Immunol. 168, 5997–6001 (2002).

14. Del Rio, L. et al. Toxoplasma gondii Triggers Myeloid Differentiation Factor 88-Dependent IL-12 and Chemokine Ligand 2 (Monocyte Chemoattractant Protein 1) Responses Using Distinct Parasite Molecules and Host Receptors. J. Immunol. 172, 6954–6960 (2004).

15. Yarovinsky, F. et al. TLR11 activation of dendritic cells by a protozoan profilin-like protein. Science 308, 1626–9 (2005).

16. Plattner, F. et al. Toxoplasma Profilin Is Essential for Host Cell Invasion and TLR11-Dependent Induction of an Interleukin-12 Response. Cell Host Microbe 3, 77–87 (2008).

17. Hitziger, N., Dellacasa, I., Albiger, B. & Barragan, A. Dissemination of Toxoplasma gondii to immunoprivileged organs and role of Toll/interleukin-1 receptor signalling for host resistance assessed by in vivo bioluminescence imaging. Cell. Microbiol. 7, 837–848 (2005).

18. Furuta, T., Kikuchi, T., Akira, S., Watanabe, N. & Yoshikawa, Y. Roles of the small intestine for induction of toll-like receptor 4-mediated innate resistance in naturally acquired murine toxoplasmosis. Int. Immunol. 18, 1655–1662 (2006).

19. Kim, L. et al. Toxoplasma gondii genotype determines MyD88-dependent signaling in infected macrophages. J. Immunol. 177, 2584–91 (2006).

20. Sukhumavasi, W. et al. TLR Adaptor MyD88 Is Essential for Pathogen Control during Oral Toxoplasma gondii Infection but Not Adaptive Immunity Induced by a Vaccine Strain of the Parasite. J. Immunol. 181, 3464–3473 (2008).

21. Mercer, H. L. et al. Toxoplasma gondii dense granule protein GRA24 drives MyD88-independent p38 MAPK activation, IL-12 production and induction of protective immunity. PLOS Pathog. (2020). doi:10.1371/journal.ppat.1008572

22. LaRosa, D. F. et al. T cell expression of MyD88 is required for resistance to Toxoplasma gondii. Proc. Natl. Acad. Sci. 105, 3855–3860 (2008).

23. Ge, Y. et al. Natural killer cell intrinsic toll-like receptor MyD88 signaling contributes to IL-12-dependent IFN-γ production by mice during infection with Toxoplasma gondii. Int. J. Parasitol. 44, 475–484 (2014).

24. Dinarello, C. A. Introduction to the interleukin-1 family of cytokines and receptors: Drivers of innate inflammation and acquired immunity. Immunological Reviews (2018). doi:10.1111/imr.12624

25. Powell, N. et al. The Transcription Factor T-bet Regulates Intestinal Inflammation Mediated by Interleukin-7 Receptor+ Innate Lymphoid Cells. Immunity 37, 674–684 (2012).

26. Monticelli, L. A. et al. Innate lymphoid cells promote lung-tissue homeostasis after infection with influenza virus. Nat. Immunol. 12, 1045–1054 (2011).

27. Klose, C. S. N. & Artis, D. Innate lymphoid cells as regulators of immunity, inflammation and tissue homeostasis. Nature Immunology 17, 765–774 (2016).

28. Abt, M. C. et al. Innate immune defenses mediated by two ilc subsets are critical for protection against acute clostridium difficile infection. Cell Host Microbe 18, 27–37 (2015).

29. Bancroft, G. J., Schreiber, R. D. & Unanue, E. R. Natural Immunity: A T-Cell-Independent Pathway of Macrophage Activation, Defined in the scid Mouse. Immunol. Rev. 124, 5–24 (1991).

30. Tripp, C. S., Wolf, S. F. & Unanue, E. R. Interleukin 12 and tumor necrosis factor α are costimulators of interferon γ production by natural killer cells in severe combined immunodeficiency mice with listeriosis, and interleukin 10 is a physiologic antagonist. Proc. Natl. Acad. Sci. U. S. A. 90, 3725–3729 (1993).

31. Hunter, C. A., Abrams, J. S., Beaman, M. H. & Remington, J. S. Cytokine mRNA in the central nervous system of SCID mice infected with Toxoplasma gondii: importance of T-cell-independent regulation of resistance to T. gondii. Infect. Immun. 61, 4038–44 (1993).

32. Park, E. et al. Toxoplasma gondii infection drives conversion of NK cells into ILC1-like cells. Elife 8, (2019).

33. Weizman, O.-E. El et al. ILC1 Confer Early Host Protection at Initial Sites of Viral Infection. Cell 171, 795–808.e12 (2017).

34. Hunter, C. A., Chizzonite, R. & Remington, J. S. IL-1 beta is required for IL-12 to induce production of IFN-gamma by NK cells. A role for IL-1 beta in the T cell-independent mechanism of resistance against intracellular pathogens. J. Immunol. 155, 4347–54 (1995).

35. Cai, G., Kastelein, R. & Hunter, C. A. Interleukin-18 (IL-18) enhances innate IL-12-mediated resistance to Toxoplasma gondii. Infect. Immun. 68, 6932–6938 (2000).

36. Kearley, J. et al. Cigarette smoke silences innate lymphoid cell function and facilitates an exacerbated type I interleukin-33-dependent response to infection. Immunity 42, 566–579 (2015).

37. Melchor, S. J. et al. IL-1R Regulates Disease Tolerance and Cachexia in Toxoplasma gondii Infection. J. Immunol. 204, 3329–3338 (2020).

38. Yap, G. S., Ortmann, R., Shevach, E. & Sher, A. A Heritable Defect in IL-12 Signaling in B10.Q/J Mice. II. Effect on Acute Resistance to Toxoplasma gondii and Rescue by IL-18 Treatment. J. Immunol. 166, 5720–5725 (2001).

39. Vossenkämper, A. et al. Both IL-12 and IL-18 contribute to small intestinal Th1-type immunopathology following oral infection with *Toxoplasma gondii*, but IL-12 is dominant over IL-18 in parasite control. Eur. J. Immunol. 34, 3197–3207 (2004).

40. Ewald, S. E., Chavarria-Smith, J. & Boothroyd, J. C. NLRP1 Is an Inflammasome Sensor for Toxoplasma gondii. Infect. Immun. 82, 460–468 (2014).

41. Gorfu, G. et al. Dual Role for Inflammasome Sensors NLRP1 and NLRP3 in Murine Resistance to Toxoplasma gondii. MBio 5, e01117–13 (2014).

42. Schmitz, J. et al. IL-33, an interleukin-1-like cytokine that signals via the IL-1 receptor-related protein ST2 and induces T helper type 2-associated cytokines. Immunity 23, 479–490 (2005).

43. Moussion, C., Ortega, N. & Girard, J. P. The IL-1-like cytokine IL-33 is constitutively expressed in the nucleus of endothelial cells and epithelial cells in vivo: A novel ‘Alarmin’? PLoS One 3, 1–8 (2008).

44. Liew, F. Y., Girard, J.-P. P. & Turnquist, H. R. Interleukin-33 in health and disease. Nat. Rev. Immunol. 16, 676–689 (2016).

45. Rostan, O. et al. Crucial and diverse role of the interleukin-33/ST2 axis in infectious diseases. Infect. Immun. 83, 1738–1748 (2015).

46. Kouzaki, H. et al. The Danger Signal, Extracellular ATP, Is a Sensor for an Airborne Allergen and Triggers IL-33 Release and Innate Th2-Type Responses. J. Immunol. 186, 4375–4387 (2011).

47. Kakkar, R., Hei, H., Dobner, S. & Lee, R. T. Interleukin 33 as a mechanically responsive cytokine secreted by living cells. J. Biol. Chem. 287, 6941–8 (2012).

48. Cohen, E. S. et al. Oxidation of the alarmin IL-33 regulates ST2-dependent inflammation. Nat. Commun. 6, 8327 (2015).

49. Humphreys, N. E., Xu, D., Hepworth, M. R., Liew, F. Y. & Grencis, R. K. IL-33, a Potent Inducer of Adaptive Immunity to Intestinal Nematodes. J. Immunol. 180, 2443–2449 (2008).

50. Silver, J. S. et al. Inflammatory triggers associated with exacerbations of COPD orchestrate plasticity of group 2 innate lymphoid cells in the lungs. Nat. Immunol. 17, 626–635 (2016).

51. Osbourn, M. et al. HpARI Protein Secreted by a Helminth Parasite Suppresses Interleukin-33. Immunity 47, 739–751.e5 (2017).

52. Ricardo-Gonzalez, R. R. et al. Tissue signals imprint ILC2 identity with anticipatory function. Nat. Immunol. 19, 1093–1099 (2018).

53. Molofsky, A. B., Savage, A. K. & Locksley, R. M. Interleukin-33 in Tissue Homeostasis, Injury, and Inflammation. Immunity 42, 1005–1019 (2015).

54. Matta, B. M. et al. IL-33 Is an Unconventional Alarmin That Stimulates IL-2 Secretion by Dendritic Cells To Selectively Expand IL-33R/ST2 ^+^ Regulatory T Cells. J. Immunol. 193, 4010–4020 (2014).

55. Schiering, C. et al. The alarmin IL-33 promotes regulatory T-cell function in the intestine. Nature 513, 564–568 (2014).

56. Spallanzani, R. G. et al. Distinct immunocyte-promoting and adipocyte-generating stromal components coordinate adipose tissue immune and metabolic tenors. Sci. Immunol. 4, eaaw3658 (2019).

57. Ito, M. et al. Brain regulatory T cells suppress astrogliosis and potentiate neurological recovery. Nature 565, 246–250 (2019).

58. Franca, R. O. R. F. O. et al. IL-33 signaling is essential to attenuate viral-induced encephalitis development by downregulating iNOS expression in the central nervous system. J. Neuroinflammation 13, 159 (2016).

59. Milovanovic, M. et al. Deletion of IL-33R (ST2) Abrogates Resistance to EAE in BALB/C Mice by Enhancing Polarization of APC to Inflammatory Phenotype. PLoS One 7, 1–13 (2012).

60. Xiao, Y. et al. Interleukin-33 deficiency exacerbated experimental autoimmune encephalomyelitis with an influence on immune cells and glia cells. Mol. Immunol. 101, 550–563 (2018).

61. Rostan, O. et al. The IL-33/ST2 Axis Is Associated with Human Visceral Leishmaniasis and Suppresses Th1 Responses in the Livers of BALB/c Mice Infected with Leishmania donovani. MBio 4, e00383–13 (2013).

62. Nabekura, T., Girard, J.-P. & Lanier, L. L. IL-33 receptor ST2 amplifies the expansion of NK cells and enhances host defense during mouse cytomegalovirus infection. J. Immunol. 194, 5948–52 (2015).

63. Bonilla, W. V. et al. The Alarmin Interleukin-33 Drives Protective Antiviral CD8+ T Cell Responses. Science (80-.). 335, 984–989 (2012).

64. Baumann, C. et al. T-bet– and STAT4–dependent IL-33 receptor expression directly promotes antiviral Th1 cell responses. Proc. Natl. Acad. Sci. U. S. A. 112, 4056–4061 (2015).

65. Still, K. M. et al. Astrocytes promote a protective immune response to brain Toxoplasma gondii infection via IL-33-ST2 signaling. PLOS Pathog. 16, e1009027 (2020).

66. Jones, L. A. et al. IL-33 receptor (T1/ST2) signalling is necessary to prevent the development of encephalitis in mice infected with Toxoplasma gondii. Eur. J. Immunol. 40, 426–436 (2010).

67. Konradt, C. et al. Endothelial cells are a replicative niche for entry of Toxoplasma gondii to the central nervous system. Nat. Microbiol. 1, (2016).

68. Van Grol, J., Muniz-Feliciano, L., Portillo, J. A. C., Bonilha, V. L. & Subaustea, C. S. CD40 induces anti-toxoplasma gondii activity in nonhematopoietic cells dependent on autophagy proteins. Infect. Immun. 81, 2002–2011 (2013).

69. Betancourt, E. D. et al. From entry to early dissemination-Toxoplasma gondii’sInitial Encounter with Its Host. Frontiers in Cellular and Infection Microbiology 9, 46 (2019).

70. Luu, L. et al. An open-format enteroid culture system for interrogation of interactions between Toxoplasma gondii and the intestinal epithelium. Front. Cell. Infect. Microbiol. 9, (2019).

71. Ju, C.-H., Chockalingam, A. & Leifer, C. A. Early Response of Mucosal Epithelial Cells during Toxoplasma gondii Infection. J. Immunol. 183, 7420–7427 (2009).

72. Hauser Jr., W. E., Sharma, S. D. & Remington, J. S. Augmentation of NK cell activity by soluble and particulate fractions of Toxoplasma gondii. J Immunol (1983).

73. Hunter, C. A. et al. The role of the CD28/B7 interaction in the regulation of NK cell responses during infection with Toxoplasma gondii. J. Immunol. 158, 2285–93 (1997).

74. Wherry, J. C., Schreiber, R. D. & Unanue, E. R. Regulation of gamma interferon production by natural killer cells in scid mice: roles of tumor necrosis factor and bacterial stimuli. Infect. Immun. 59, 1709–15 (1991).

75. Johnston, L. K. et al. IL-33 Precedes IL-5 in Regulating Eosinophil Commitment and Is Required for Eosinophil Homeostasis. J. Immunol. 197, 3445–3453 (2016).

76. Christian, D. A. et al. cDC1 Coordinate Innate and Adaptive Responses in the Omentum required for T cell Priming and Memory. bioRxiv 2020.07.21.214809 (2020). doi:10.1101/2020.07.21.214809

77. Jackson-Jones, L. H. et al. Fat-associated lymphoid clusters control local IgM secretion during pleural infection and lung inflammation. Nat. Commun. 7, 12651 (2016).

78. Buscher, K. et al. Protection from septic peritonitis by rapid neutrophil recruitment through omental high endothelial venules. Nat. Commun. 7, 1–7 (2016).

79. Monticelli, L. A. et al. IL-33 promotes an innate immune pathway of intestinal tissue protection dependent on amphiregulin–EGFR interactions. Proc. Natl. Acad. Sci. 112, 10762–10767 (2015).

80. Gazzinelli, R. T. et al. Interleukin 12 is required for the T-lymphocyte-independent induction of interferon y by an intracellular parasite and induces resistance in T-cell-deficient hosts (Toxoplasma gondii/natural killer cells). Immunology 90, 6115–6119 (1993).

81. Hunter, C. A., Candolfi, E., Subauste, C., Van Cleave, V. & Remington, J. S. Studies on the role of interleukin-12 in acute murine toxoplasmosis. Immunology 84, 16–20 (1995).

82. Yap, G., Pesin, M. & Sher, A. Cutting Edge: IL-12 Is Required for the Maintenance of IFN-γ Production in T Cells Mediating Chronic Resistance to the Intracellular Pathogen, Toxoplasma gondii. J. Immunol. 165, 628–631 (2000).

83. Wilson, D. C., Matthews, S. & Yap, G. S. IL-12 Signaling Drives CD8 + T Cell IFN-γ Production and Differentiation of KLRG1 + Effector Subpopulations during Toxoplasma gondii Infection. J. Immunol. 180, 5935–5945 (2008).

84. Mordue, D. G., Monroy, F., La Regina, M., Dinarello, C. A. & Sibley, L. D. Acute Toxoplasmosis Leads to Lethal Overproduction of Th1 Cytokines. J. Immunol. 167, 4574–4584 (2001).

85. Struck, D. et al. Treatment with interleukin-18 binding protein ameliorates Toxoplasma gondii-induced small intestinal pathology that is induced by bone marrow cell-derived interleukin-18. Eur. J. Microbiol. Immunol. 2, 249–257 (2012).

86. Muñoz, M. et al. Interleukin-22 Induces Interleukin-18 Expression from Epithelial Cells during Intestinal Infection. Immunity 42, 321–331 (2015).

87. Villeret, B. et al. Blockade of IL-1R signaling diminishes Paneth cell depletion and Toxoplasma gondii induced ileitis in mice. Am. J. Clin. Exp. Immunol. 2, 107–16 (2013).

88. Walker, W., Aste-Amezaga, M., Kastelein, R. A., Trinchieri, G. & Hunter, C. A. IL-18 and CD28 use distinct molecular mechanisms to enhance NK cell production of IL-12-induced IFN-gamma. J. Immunol. 162, 5894–901 (1999).

89. Man, S. M. & Kanneganti, T.-D. Regulation of inflammasome activation. Immunol. Rev. 265, 6–21 (2015).

90. Zediak, V. P. & Hunter, C. A. IL-10 fails to inhibit the production of IL-18 in response to inflammatory stimuli. Cytokine 21, 84–90 (2003).

91. Lima, T. S., Gov, L. & Lodoen, M. B. Evasion of human neutrophil-mediated host defense during Toxoplasma gondii infection. MBio 9, (2018).

92. Liu, Y. et al. Toxoplasma gondii Cathepsin C1 inhibits NF-κB signalling through the positive regulation of the HIF-1α/EPO axis. Acta Trop. 195, 35–43 (2019).

93. DeLaney, A. A. et al. Caspase-8 promotes c-Rel–dependent inflammatory cytokine expression and resistance against Toxoplasma gondii. Proc. Natl. Acad. Sci. U. S. A. 116, 11926–11935 (2019).

94. Sandstrom, A. et al. Functional degradation: A mechanism of NLRP1 inflammasome activation by diverse pathogen enzymes. Science (80-.). 364, (2019).

95. Snelgrove, R. J. et al. Alternaria-derived serine protease activity drives IL-33-mediated asthma exacerbations. J. Allergy Clin. Immunol. 134, (2014).

96. Chen, W.-Y., Hong, J., Gannon, J., Kakkar, R. & Lee, R. T. Myocardial pressure overload induces systemic inflammation through endothelial cell IL-33. Proc. Natl. Acad. Sci. U. S. A. 112, 7249–7254 (2015).

97. Tashiro, H. et al. Interleukin-33 from Monocytes Recruited to the Lung Contributes to House Dust Mite-Induced Airway Inflammation in a Mouse Model. PLoS One 11, e0157571 (2016).

98. Li, C. et al. Interleukin-33 Increases Antibacterial Defense by Activation of Inducible Nitric Oxide Synthase in Skin. PLoS Pathog. 10, e1003918 (2014).

99. Reverchon, F. et al. IL-33 receptor ST2 regulates the cognitive impairments associated with experimental cerebral malaria. PLoS Pathog. (2017). doi:10.1371/journal.ppat.1006322

